# State-dependent modulation of optic flow-processing lobula plate cells in butterflies

**DOI:** 10.64898/2026.09.23.749361

**Authors:** Alexandra M. Yarger, Jack A. Supple, Holger G. Krapp

**Affiliations:** Department of Bioengineering, Imperial College London, London SW7 2AZ, United Kingdom

## Abstract

Increasing experimental evidence suggests that biological systems cancel predictable components of sensory signals while maintaining sensitivity to externally induced state changes. This strategy provides task-specific sensor responses for posture, locomotion, and gaze control. A prime example is found in interneurons that respond to visual image shifts resulting from the relative motion between an animal’s eyes and its visual surroundings. Such optic flow-processing interneurons, found across phyla and are particularly well characterized in Dipteran and other flying insects. We studied optic flow-sensitive interneurons in the optic lobe of the Monarch butterfly, whose large and highly contrasted wings sweep through the visual field with every wing-beat cycle, potentially obscuring interneuron output signals. Our results show baseline spiking activity increases when animals flap their wings, and individual spikes are phase-locked to the wing-beat cycle, even in the dark, when no visual motion input is available. A qualitative estimate of interneuron responses to directional wing motion through their receptive field is not sufficient to explain the recorded activity patterns. Our results suggest that an additional internal signal suppresses responses to wing-induced visual motion to support effective vision-based stabilization reflexes. These findings support the principle that self-generated signals are suppressed while sensitivity to external modulation is preserved.

## Introduction

The visual motion patterns experienced during self-movement are used across phyla and in autonomous robotics for state estimation and to control stabilization reflexes. Directionally selective visual neurons that are tuned to body rotations and translations have been reported in different species from insects to primates^1–3^. The panoramic image shift experienced as a result of the relative movement between the observer and the visual scene is called optic flow^4^. Directionally selective neurons are tuned to respond most strongly to particular types of motion patterns – or optic flow fields – and provide vital information for postural control and gaze stabilization^5,6^.

However, a fundamental challenge for such systems is distinguishing the sensory consequences of self-motion (re-afference), and those arising from external changes (ex-afference) - a failure to do so would result in a futile cycle of responding to one’s own motion^7^. The nervous system is thought to overcome this re-afferent reflex trap by constructing an internal model of the body and its configuration within the world^8^. By running copies of motor commands (‘efference’ copies) through this internal ‘forward’ model, the brain can simulate the sensory consequences of volitional self-motion and prepare sensory systems to cancel anticipated re-afference^8^.

In flies, motor-related signals akin to efference copies have been reported in optic flow-sensitive interneurons in the lobula plate^9–12^. These lobula plate tangential cells (LPTCs) are sensitive to specific patterns of optic flow associated with rotations of the animal^13^. During voluntary flight turns, LPTCs receive motor-related inputs that cancel the visual responses to the resulting self-generated optic flow, thereby suppressing visual re-afference that would otherwise drive compensatory stabilisation reflexes^9–12^. Such predictive modulation is therefore consistent with the use of an internal model of how the animal’s own movements are expected to transform the visual scene.

But what happens when an observer’s own body parts move through the receptive field of an optic-flow processing neuron? If appendage motion is sufficient to evoke a visual response, this could also disrupt the stable and accurate monitoring of global self-motion and would therefore benefit from predictive cancellation of re-afference. However, such predictive cancellation would require knowledge of how the body is configured and how appendage movements project onto the retina.

Lepidopterans like the Monarch butterfly (*Danaus Plexippus*) are particularly susceptible to this problem. Their wings are so large, that with every wingstroke, a significant portion of their visual field is occluded. This not only prevents them from seeing past their wings but could also potentially activate the same visual neurons that they rely on to detect wide-field motion and maintain a stable gaze and attitude during flight. Here, we map the receptive field organization of wide-field sensitive neurons in the lobula plate of Monarch butterflies and characterize their state-dependent modulation during flapping.

## Results

### Receptive field organization

We characterized the receptive field organization of 212 motion-sensitive visual interneurons in *D. Plexippus.* Most of these interneurons feature large receptive fields that cover extended parts of both visual hemispheres. They are located in the lobula plate, an output stage of the animals visuomotor pathway, and are accordingly referred to as Lobula Plate Cells (LPCs). To determine the cells’ local motion sensitivity (LMS) and local preferred direction (LPD), we mounted butterflies on a goniometric recording platform (GRP; Fig. 1A) and recorded extracellularly their spiking activity upon visual motion stimulation. The GRP rotated the butterfly around two orthogonal axes in front of a stationary monitor displaying a small circular patch of black-and-white gratings moving sequentially in eight directions. By positioning this stimulus at different visual azimuths and elevations, we sampled local motion responses across the visual field (Fig. 1B). At each location, the LMS and LPD together defined a local response vector describing directional motion sensitivity (Fig. 1C). When plotted as a function of azimuth and elevation in a 2D cylindrical projection, these vectors reveal the spatial organisation of the cell’s receptive field (Fig. 2A). Cells with incoherent response vector orientations (Fig.S1A), target-selective cells (Fig. S1B), and translation sensitive cells (Fig. S1C) were not included in further analysis, leaving a total of 70 wide-field rotation sensitive LPCs.

**Figure 1:**
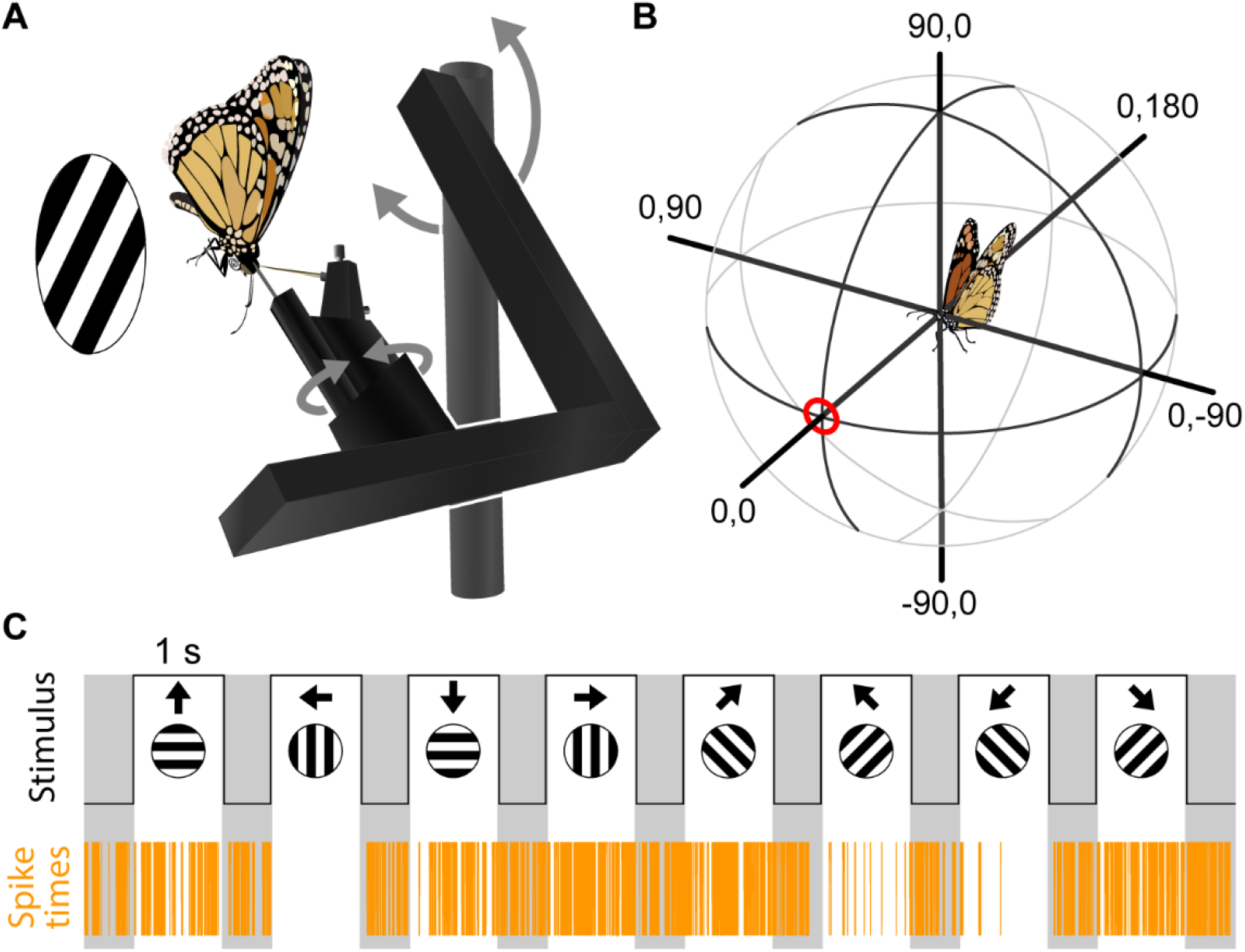
**A)** Experimental setup: Goniometric Recording Platform. **B**) The centre of the butterfly head defines the origin of a spherical coordinate system where numbers indicate azimuth and elevation. 0° azimuth, 0° elevation indicates the animal’s forward direction. **C**) Moving grating stimuli (top) to determine local motion sensitivity and preferred direction based on the spiking activity of the LPC (bottom). Grey regions indicate no stimulus (dark) periods.

**Figure 2:**
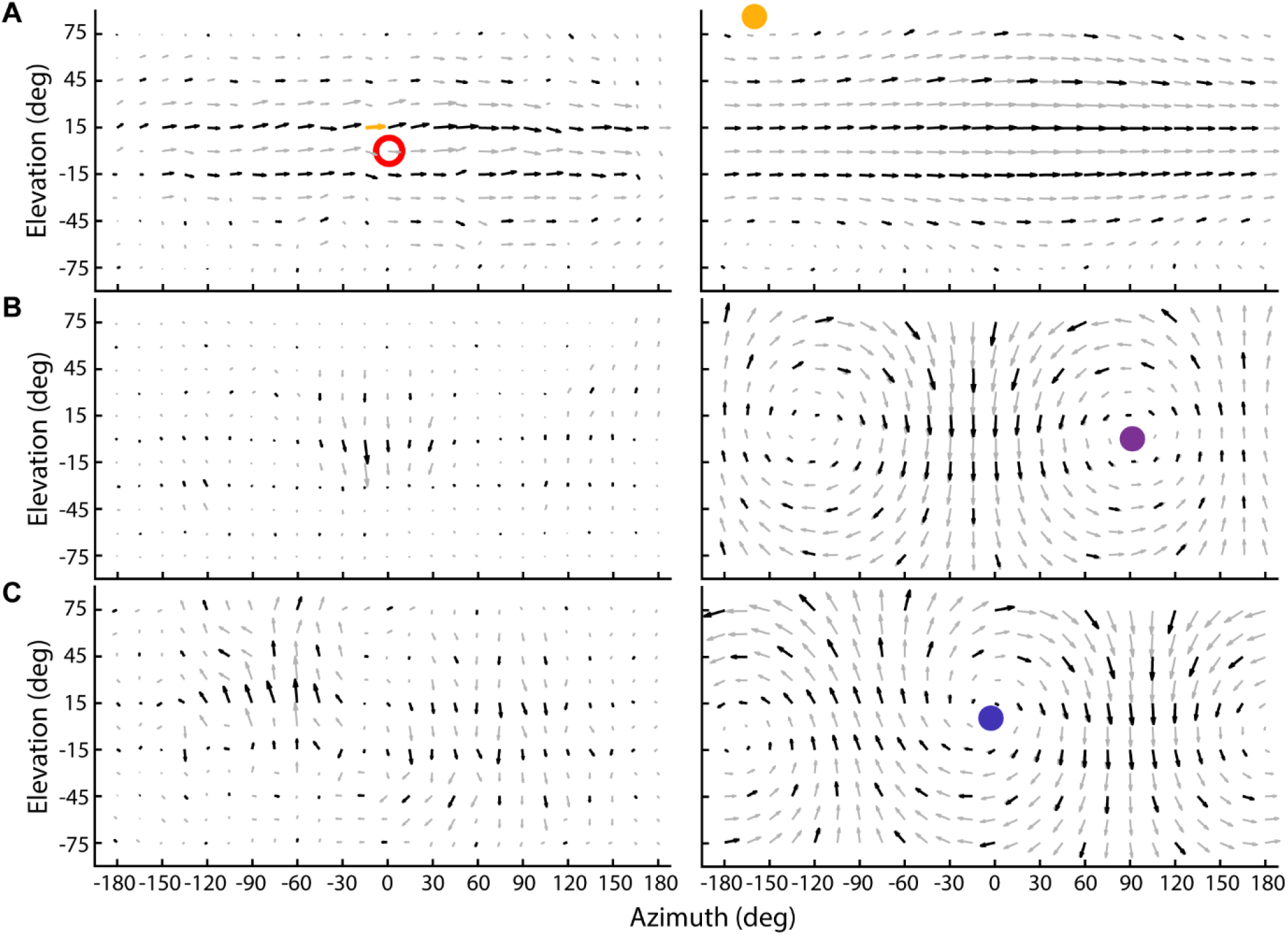
**A)** Example receptive field (left) of a yaw-sensitive cell. The red circle indicates the direction the animal is facing. The yellow vector is based on the recording shown in Fig. 1C. Black vectors show measured data; grey vectors have been interpolated. The Koenderink– van Doorn (KvD) algorithm was applied to the measured data to obtain the preferred self-motion parameters that define optic-flow fields, the cells are tuned to detect. The optic flow fields including the orientation of the rotation axis (coloured dots) are plotted on the right. **B**) Example of a pitch-sensitive cell. **C**) Example of a roll-sensitive cell.

Figure 2 (left column) shows example receptive fields of LPCs classified as wide-field cells. Qualitatively, they are tuned to respond to optic flow-field induced by yaw- (Fig 2A), pitch- (Fig 2B), and roll- (Fig 2C) rotations. All cells have a binocular receptive field organization with a coherent arrangement of response vectors reminiscent of a rotational flow field structure. To quantitatively assess the self-motion parameters these wide-field LPC receptive fields, we applied an iterative least-square algorithm proposed by Koenderink–van Doorn (KvD; 1987) and plotted the resulting optic flow fields next to the corresponding receptive fields (Fig. 2, right column). The location of the preferred rotation axis was used to classify each cell into one of three categories.

To visualize the results of all 70 LPCs studied, we plotted each cell’s preferred axis of rotation in a cylindrical projection (Fig. 3A) and a spherical representation of the animal’s visual field (Fig. 3B). Each dot and arrow in the cylindrical projection and spherical representation, respectively, corresponds to the body axis about which the butterfly would need to rotate clockwise to strongly excite that cell. All recordings were obtained from the right side of the brain. To account for the cells’ corresponding contralateral counterparts in the left side of the brain we also plotted each cell’s mirror transformed preferred axis of rotation.

**Figure 3:**
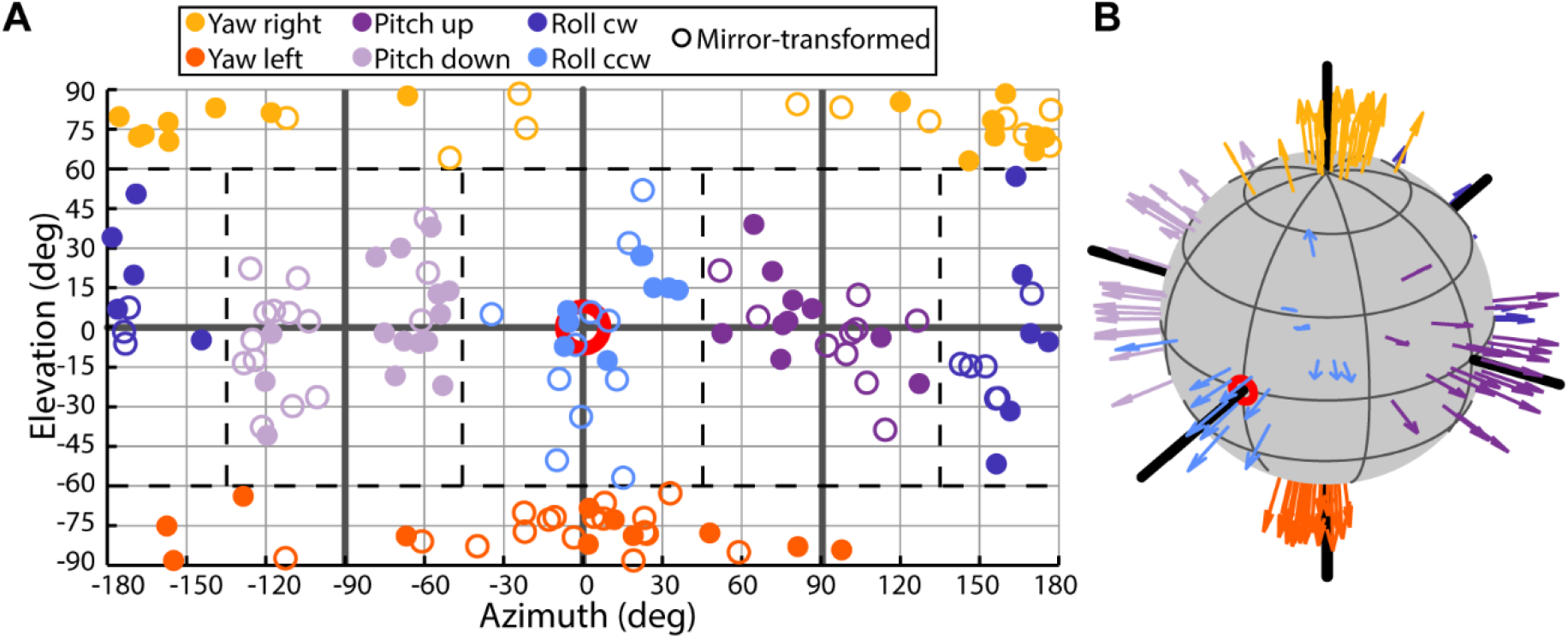
**A)** Preferred axes of rotation for all recorded cells (dots) and their mirror-transformed counterparts (empty circles) plotted in a cylindrical projection as a function of azimuth and elevation. Colour indicates cell type. Dotted lines represent cutoff regions for cell classifications. **B**) Preferred axes of rotation plotted in a spherical representation to emphasize the tight clustering of yaw-sensitive LPCs in the pole region of the visual field.

### Receptive field variability

An increased level of variability – i.e. a deviation from a coherent distribution of local response vectors within the measured receptive field (Fig. 4A) – was frequently observed as a result of unexpected bursts in spiking activity. These bursts often correlated with wing movements. Restraining the wings did not significantly reduce variability, but removing the wings and restraining the thorax with beeswax did significantly reduce the angular variance of the preferred direction across trials (Fig. 4B; N=19 animals).

**Figure 4:**
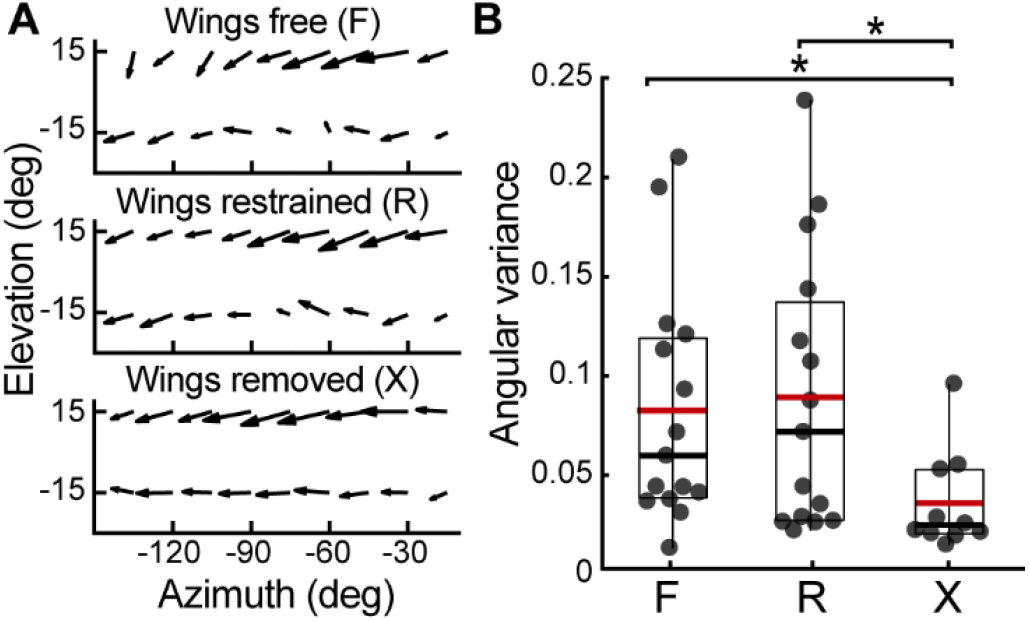
**A)** Measured preferred directions when wings were free to move (top), restrained (middle), or removed (bottom). **B**) Angular variance of yaw cells measured from the same region of the receptive field (Azimuth -150 to 0, Elevation -30 to 30) in animals with wings free (F), restrained (R), or removed (X). Removing the wings significantly decreased angular variance (N=19 LPCs, 19 animals; p = 0.04, 0.03).

### Visual feedback from high-contrast wings

To further investigate whether the observed changes in LPC activity were a result of visual or mechanosensory feedback from the wings, we recorded LPCs during flapping in the light and dark without any external visual stimulus. There were no significant differences in spike rate between the light and dark conditions, but there was a significant increase in spike rate during flapping (Fig. 5A-B). On average, the spike rate increased by 21.4 ± 5.81 Hz from baseline during flapping.

**Figure 5:**
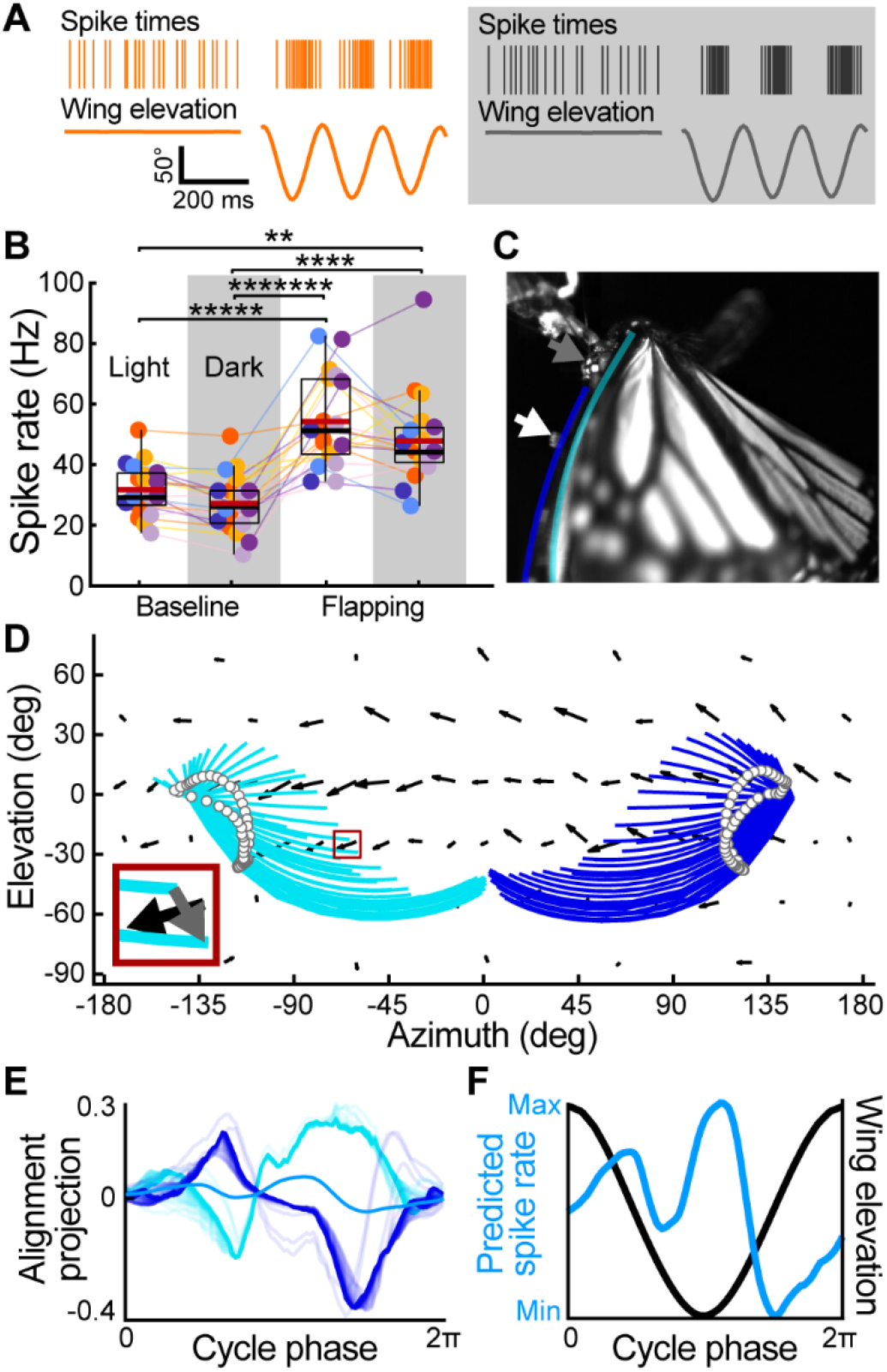
**A)** Example flapping bouts. Spike times (top) and wing elevation (bottom) recorded in the light (orange) and the dark (black). **B**) Butterfly LPC baseline spike rate compared with flapping spike rate in the light and dark (N=30 LPCs, 24 animals). Grey background indicates activities recorded in the dark. The spike rate is significantly higher and more variable during flapping. Light condition has no effect on spike rate. **C**) Downstroke of wingbeat cycle showing wings occlude the visual field. Grey arrow indicates head. White arrow indicates location of wing marker. **D**) Wing occlusion of the visual field during a single wingbeat cycle. Black arrows show local response vectors of the LPC, the spiking activity of which is shown in panel A. White dots represent tracked wing marker. Cyan and dark blue lines represent the leading edge of the left and right wings respectively. The inset shows an example of the movement of the wing tip between two consecutive frames (grey arrow) and the LPC’s response vector (black arrow) at the same location of the visual field where the wing movement occurred. **E**) Averaged dot products between LPC response vectors and wing movements along the leading edge for the left (cyan), right (dark blue), and both (light blue) wings. **F**) Normalized wing elevation (black) and predicted spike rate averaged across both wings and all wingbeats (light blue).

Butterfly wings are large enough that, at the bottom of the downstroke, the wings cover more than half of their eyes (Fig. 5C). The sweeping motion of this high contrast structure across the visual field is very similar to the black and white gratings used to stimulate LPCs. Flapping wings could therefor result in activity changes in LPCs sensitive to rotation-induced wide field motion. We tested whether visual motion caused by the wings sweeping across a cell’s receptive field could predict a cell’s spiking response during flapping. To this end, we measured the relative speed and direction of 100 points along the leading edge of the wing and projected the resulting vectors into the cell’s local response vectors at corresponding locations in the visual field (Fig. 5D). By averaging the local dot products (Fig. 5E) across both wings for 30 wingbeats we estimated the predicted spike rate throughout the wingbeat cycle (Fig. 5F).

### Phase-locking with wingbeat cycle

To determine whether the spike rate predicted from the position of the wing during flapping matched the cell’s actual activity, we measured the spike count and instantaneous firing frequency (Fig. 6A) for the same 30 wingbeats used to calculate the predicted response. We found that the expected response based solely on wing movement did not predict the measured spiking activity (Fig. 6A) for all cell types (Fig. S2). This mismatch between the actual spike rate and the expected response suggests that the modulated LPC activity cannot be explained by the visual motion of the wing alone. Further, phase locking with wing elevation was also observed in the dark for all cell types. There were also no significant differences in the phase locking strength or timing under light and dark conditions (Fig. 6A-B) for all yaw type cells (N=6 animals, 8 cells). – i.e. cells that responded to horizontal motion. This finding confirmed that phase locking in yaw-sensitive LPCs is driven by non-visual inputs. However, pitch and roll type cells – i.e. cells that responded to vertical motion – showed significantly weaker phase locking in the dark which was also shifted in the absence of visual feedback (Fig. 6C-D; N=6 animals, 8 cells). This indicates that visual information due to wing movement may still modulate the activity of these cells.

**Figure 6:**
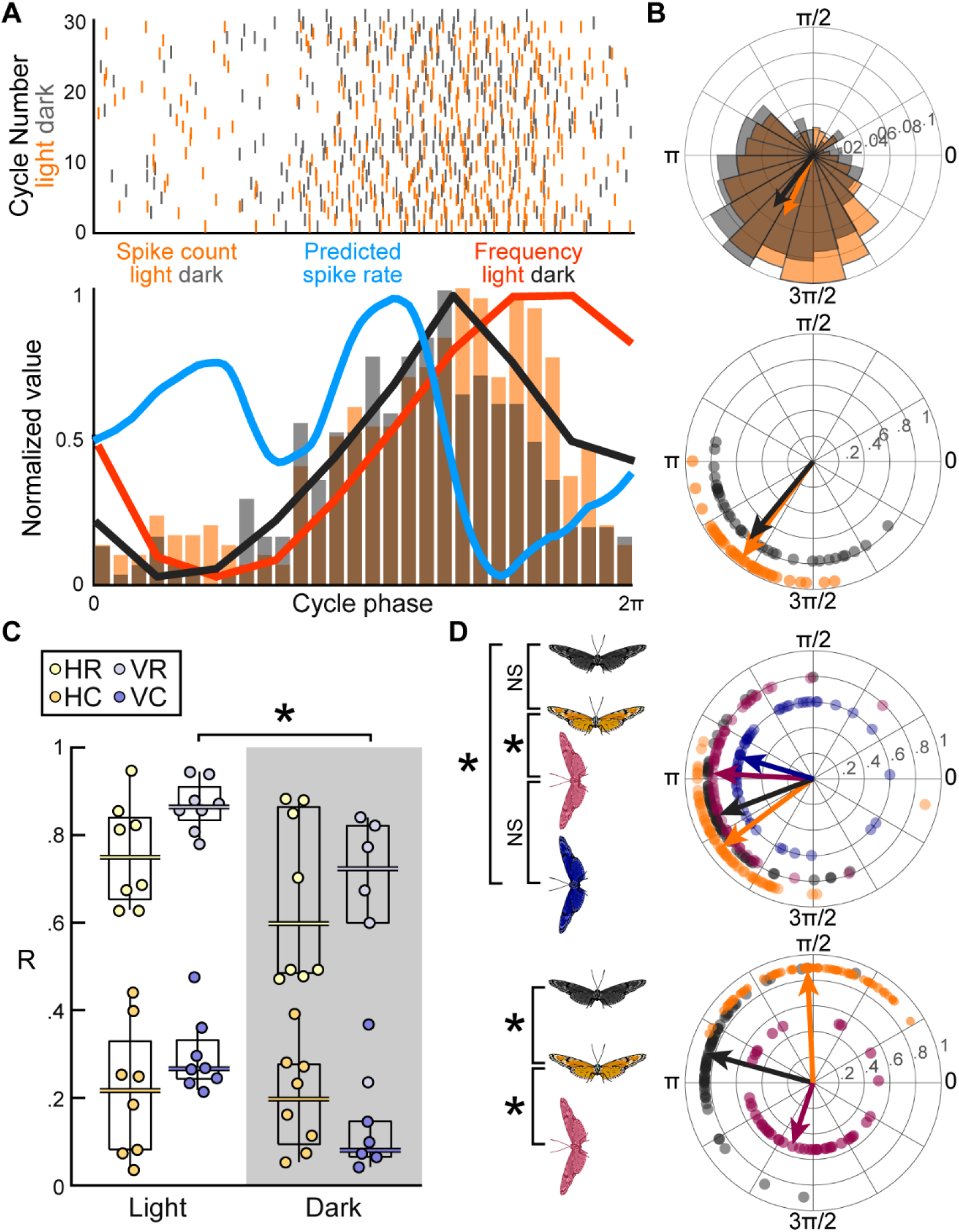
**A**) Raster plot of spike times (top panel) relative to wingbeat phase in the light (orange) and in the dark (grey). Predicted spike response (blue line; see Fig. 5F), binned spike counts (orange & grey histograms), and average instantaneous spike frequency in the light and dark (orange and black lines) all normalized to wingbeat cycle. **B**) Polar plots showing phase locking pattern of spikes relative to the wingbeat cycle in the light (orange) and dark (grey). Top panel uses binned spike counts. Bottom panel uses instantaneous spike frequency. **C**) Phase locking magnitude calculated from instantaneous spike rate (R) and spike count (C) is significantly stronger in the light for cells that respond to vertical motion (V), but not for cells that respond to horizontal motion (H). **D**) Example polar plots for yaw (top) and pitch (bottom) cells recorded during flapping in the light (orange) and the dark (grey) in upright and sideways (pink and purple) orientations. Body orientation significantly changed phase of activation across all cells, but light condition only affected the phase of vertical cells.

### State-dependent modulation

Cells with very similar receptive fields did not always exhibit the same phase of activation relative to the wingbeat cycle (Fig S3). This could be explained by individual differences in morphology and tuning if the phase timing within individuals always occurred at a fixed point of the wingbeat cycle, remaining constant regardless of the overall trajectory of the wing. Alternatively, the phase of activation could also be affected by the wing’s kinematics (e.g. asymmetry between the left and right wings). To determine whether phase timing is modulated by wing trajectory, we rotated the butterflies 90° and recorded LPC activity during flapping. Rotations resulted in asymmetry between the left- and right-wing trajectories. This occurs because the butterflies attempt to correct their orientation by increasing the maximum and minimum angle of elevation for one wing and decreasing them for the other. We found that, for all cell types, the phase of activation was significantly different for upright and rotated animals (Fig. 6D, Fig. S3; N=6 animals, 6 cells).

The light condition did not change the phase of activation for horizontal cells (i.e. there was no significant difference between animals rotated in the light and animals rotated in the dark). This showed that horizontal LPC activity is modulated by the wing motion itself, not by the visual stimulation resulting from the wings sweeping through their receptive field.

## Discussion

### Behavioural state modulation

The tuning of sensory neurons is matched to the functional requirements of the individual, but they can also change depending on an animal’s internal state, or environmental conditions^14–16^. For example, visual sensitivity in flies has been shown to change during walking vs. flying^17^ and when flies are hungry vs. sated^18^. LPC baseline activity increased during flapping, even in the absence of any visual input. This is consistent with findings in fly lobula plate tangential cells (LPTCs). Previous work showed that LPTC baseline activity increases when an octopamine agonist is used to mimic the flight state. This increase in baseline firing rate also increases the signalling range and the rate of information transfer^19–22^. The state-dependent modulation shown here may allow butterfly LPCs to operate in different dynamic ranges during flapping and quiescence, which could provide a functional advantage because relevant optic flow signals occupy different spatiotemporal ranges across different behaviours.

### Separating self-generated motion from optic flow

Insects have evolved specialized sensory modalities for selectively detecting task-relevant inputs with different sensory systems working together to produce complex and specialized behaviours. Flies use inertial sensors to maintain stability while chasing conspecifics at high speed on the wing^23,24^. The high acuity of dragonfly vision helps them track prey and engage in aerial combat^25^. Butterflies use the position of the sun, as well as polarized light and magnetic fields to navigate thousands of miles during migrations^26–28^. Fast mechanosensory signals enable immediate body adjustments, while slower temporal dynamics are supported by vision^29–32^ to extend the overall dynamic input range insects can handle.

Lepidopteran wings present an atypical control problem. Any self-motion generates optic flow, which provides a rich source of information to visually sense state changes. This is however based on the animal’s movement relative to its environment and may not normally be superimposed by actively moving body parts such as limbs or, in this case, coloured wings. The distinctive black and orange stripes of Monarch butterfly wings exacerbate this issue, as the LPCs responsible for detecting self-motion signals respond most strongly to precisely this type of high-contrast pattern. Yet, LPCs do not respond in a way that is consistent with what we predicted for visual stimuli that resulted from the flapping movement of the wings.

There were some obvious differences between receptive field types, which we were able to divide into two categories. Yaw cells fell into one group we called “horizontal cells” (HCs) and the remaining roll- and pitch-sensitive cells were classified as “vertical cells” (VCs). VCs are more likely to be involved in attitude control (stability responses) and HCs are likely used for spontaneous or deliberate steering manoeuvres. VCs were more weakly phase locked in the dark. This may indicate that these cells maintain some level of visual sensitivity, which could be useful for reflexive stability control.

If a butterfly had an internal representation (akin to an efference copy) of the expected wing-induced visual motion and a gust of wind altered the wing’s trajectory, the mismatch between the expected and actual visual inputs could be sent to the flight motor as a corrective control command. Robotic systems frequently use similar visual measurements of self-generated motion (visual servoing) to control their state changes^33^.

Visual stimulation due to flapping wings is less likely to activate yaw-sensitive cells, simply because the wing trajectory contains mostly vertical motion components. Even so, the remaining horizontal wing motion components, were not reflected in these LPC’s recorded activity patterns when the animal was flapping its wings. Further, there were no observable differences in HC activity in the light versus the dark. These results indicate that horizontal cells are functionally blind to the wings and primarily respond to visual inputs originating from external sources (not the wing’s motion). As stated previously, horizontal motion is associated with steering control, so the motor commands that elicit turning (and the resulting mechanosensory feedback) is likely tied to the activity of these cells that are tuned to detect turning-induced optic flow. The mismatch we see between the expected response to wing motion and the recorded HC spiking activity, in combination with the strong phase locking to the wing motion that persists in the dark, indicates that there is indeed some motor or mechanosensory feedback modulating HC activity. This modulation is similar to the efferent-copy based suppression of the self-generated motion described in flies^9–12^. Though we did not find a clear cancellation signal, internal representations do not always serve to cancel anticipated re-afference. Their effects depend on the behavioural context and circuits involved. For example, while flight-related motor signals in flies have been shown to supress visual responses to self-generated motion, walking-related motor signals can also amplify them^9–12,34^.

Butterfly LPC receptive fields are strongly binocular (Figure S6). As a result, visual inputs from the left and right wings can be additive or subtractive depending on a cell’s receptive field. For a pure yaw cell with equal sensitivity across both hemispheres, the visual motion from left and right wings flapping symmetrically would theoretically result in zero net activation. Indeed, even the averaged dot product calculated from naturally variable wing movements and the response vectors of an inhomogeneous yaw receptive field predicted partial cancellation of the overall signal (Fig. 5E). The pronounced binocularity across LPCs could therefore partially contribute to insensitivity of HCs to visual inputs resulting from wing motion.

### Internal state modulation

Phase locking persists in the dark, so it must originate from an internal signal. The most straightforward sources may be: (i) mechanosensory/proprioceptive signals from the wings. The most likely candidates being strain sensing campaniform sensilla located near the wing base or at the wing hinge^35,36^, (ii) an efference copy/motor-related signal, or (iii) a combination of (i) and (ii).

Phase locking did not consistently occur for every type of wing movement and strict criteria were necessary to separate true flapping trials from general wing movement. It is therefore unlikely that phase locking is purely a result of mechanosensory feedback. Further, in animals that were motivated to flap continuously in the dark for longer durations, we occasionally observed phase jumps, despite no measurable changes in the wing kinematics. (Fig. S7A; Vid. S1). Subtle mechanical differences, that were not visible to us in our analysis could potentially account for these phase jumps. However, the absence of any detectable kinematic changes in these cases, the variability of phase timing for similar cell types across individuals, and the measured phase changes that occurred in response to kinematic differences that *were* detectable (asymmetry as a result of rotation; Figs. 6D & S3) are all more consistent with a centrally generated signal linked to the motor command itself rather that the feedback from the wing movement. Together, these findings suggest that LPC activity is influenced more strongly by centrally generated signals associated with the motor commands that drive wing motion than by the optic-flow signals that arise as a result of the wing movement as it sweeps across the butterfly’s visual field.

## Methods

### Receptive field characterization

To characterize the receptive field properties of the LPCs we performed extracellular recordings from the lobula plate in butterflies mounted on a goniometric recording platform (Fig. 1A). Animals were rotated to various azimuths between -180 and 180 degrees and elevations from -75 to 75 (Fig. 1B). A stationary monitor was used to stimulate the eyes with black and white gratings which moved in eight different directions. We counted the number of spikes elicited by each direction of visual motion and used these values to calculate the response-weighted resultant vector. The angle of this vector represents the neuron’s preferred direction of motion, and the size represents the strength of the directional response (Fig. 1C).

Butterflies were left fully intact and free to move their wings during recordings. Restraining the wings did not reliably reduce variability because, in many cases animals continued to contract their muscles unless the wings were removed and the thorax, abdomen, wing bases, leg bases, and mouthparts were fixed in place using beeswax (Fig. 2B). Rather than subject animals to this highly invasive manipulation, maps were averaged across 2-6 repeated trials where possible (Fig. S4). Stronger responses at a particular region of the receptive field were less variable across trials, demonstrating that the preferred directions were not innately variable and noise within individual trials was not representative of the cell’s true selectivity.

### Preferred axes

To quantitatively estimate each cell’s preferred rotational and translational self-motion parameters, we applied the iterative least-square algorithm developed by Koenderink and van Doorn^37^ (KvD), to each receptive field (Fig. 2A-C). We quantified each cell’s preference for rotation relative to translation using a rotation selectivity ratio, ***S***, where ***R*** and ***T*** represent the rotation and translation vectors, respectively. ***R*** and ***T*** were obtained from the KvD to describe an optic flow field that matches best the receptive field organization of a given LPC. Only cells with *S* > 0.4 were classified as rotation-sensitive and used for further analysis.

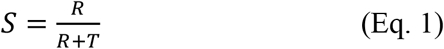

### Mirror transformation

All recordings were obtained from the right side of the brain. To account for the corresponding contralateral cell population and to approximate the full bilateral organization of the system, each receptive field was duplicated, mirror-transformed about the y-axis, and the preferred axis of rotation was re-estimated from the transformed map (Fig. 2G). This approach assumes bilateral symmetry of motion-sensitive circuitry and allows the population-level structure to be visualized more completely.

### Flapping

Flapping was induced by touching the feet or abdomen with a paintbrush. LPC activity during flapping was recorded in the dark—without visual feedback—as well as with lights on, so that butterflies could see their wings moving. 850 nm infrared light was used to illuminate the subject for high-speed videography in the dark. This wavelength is far outside the spectral sensitivity of Monarch butterflies^38^. As a baseline comparison, we also recorded segments where the animal was not moving at all (in both the light and dark). Some animals were also rotated 90° to induce asymmetry in the left- and right-wing trajectories.

### Phase locking

Spike phase was calculated relative to the wingbeat cycle. The start of the downstroke was defined as 0, and the end of the upstroke was 2π. We used a Rayleigh test to measure whether spike phases (calculated from binned spike counts) were uniformly distributed across the wingbeat cycles. As a secondary metric for phase locking, we measured the instantaneous firing frequency and used these values to calculate the mean resultant vector for each wingbeat cycle. To account for variable baseline activity, periods with higher firing frequency were more strongly weighted when calculating the mean phase. A permutation test was used measure significance of phase locking for each trial.

Many animals were not motivated to flap continuously and most flapping bouts only included 2-5 full wingbeats, which is not sufficient for phase analysis. Therefore, bouts with fewer than 8 consecutive wingbeats were excluded as well as animals that did not fully open their wings when flapping. The minimum wing elevation range of motion used was 90°.

At the start of each bout of flapping there was consistently a large burst of activity. The timing of this burst was consistent across cells (p=7 × 10^−64^; Fig. S5). This burst in activity, was excluded from further phase analyses, as it was not always consistent with the phase of activation for subsequent oscillations.

### Predicted spiking response

To estimate how LPCs would respond to the visual motion of the wing moving across the butterfly’s visual field, we filmed the wings using two Photron SA3 cameras recording at 125-500 fps. We tracked a point (*i*) on the leading edge of the wing during flapping and calculated its relative speed (*v_w_*) and direction (*θ_w_*) of motion for each frame. We compared these values to the preferred motion magnitude (*v_k_*) and direction (*θ_k_*) of the cell taken from its receptive field map, at the corresponding location of the wing marker in the visual field. Using these values, we computed the predicted spiking response 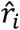, as:

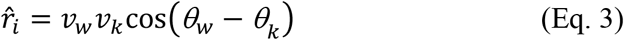

Positive values indicate that the actual motion is aligned with the preferred direction, whereas negative values indicate that the motion is opposite to the preferred direction. To estimate the contribution of the entire leading edge, we projected a line from the wing base, through the tracked point, to the wing tip in the 3D coordinate system. For each frame, this line was then plotted on the 2D cylindrical projection and used to repeat the computation for 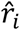. We did this for 100 points along this line to represent the leading edge of the wing). Here, *i* represents each of the 100 points along the leading edge, and the combined predicted spiking response 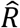, is:

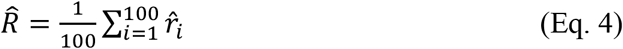

### Binocularity

To calculate the binocularity ratio, *B*, we calculated the ratio between the mean motion sensitivity in the left, *S_L_*, and right, *S_R_*, hemispheres. Values closer to 1 indicate high binocularity, whereas values near 0 reflect predominantly monocular responses.

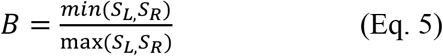

## Acknowledgements

This work was financially supported by the Air Force Office of Scientific Research (AFOSR; grant FA8655-23-1-7049 to HGK).

## Funder Information Declared

United States Air Force Office of Scientific Research, https://ror.org/011e9bt93, FA8655-23-1-7049

## Supplemental Figures

**Figure S1:**
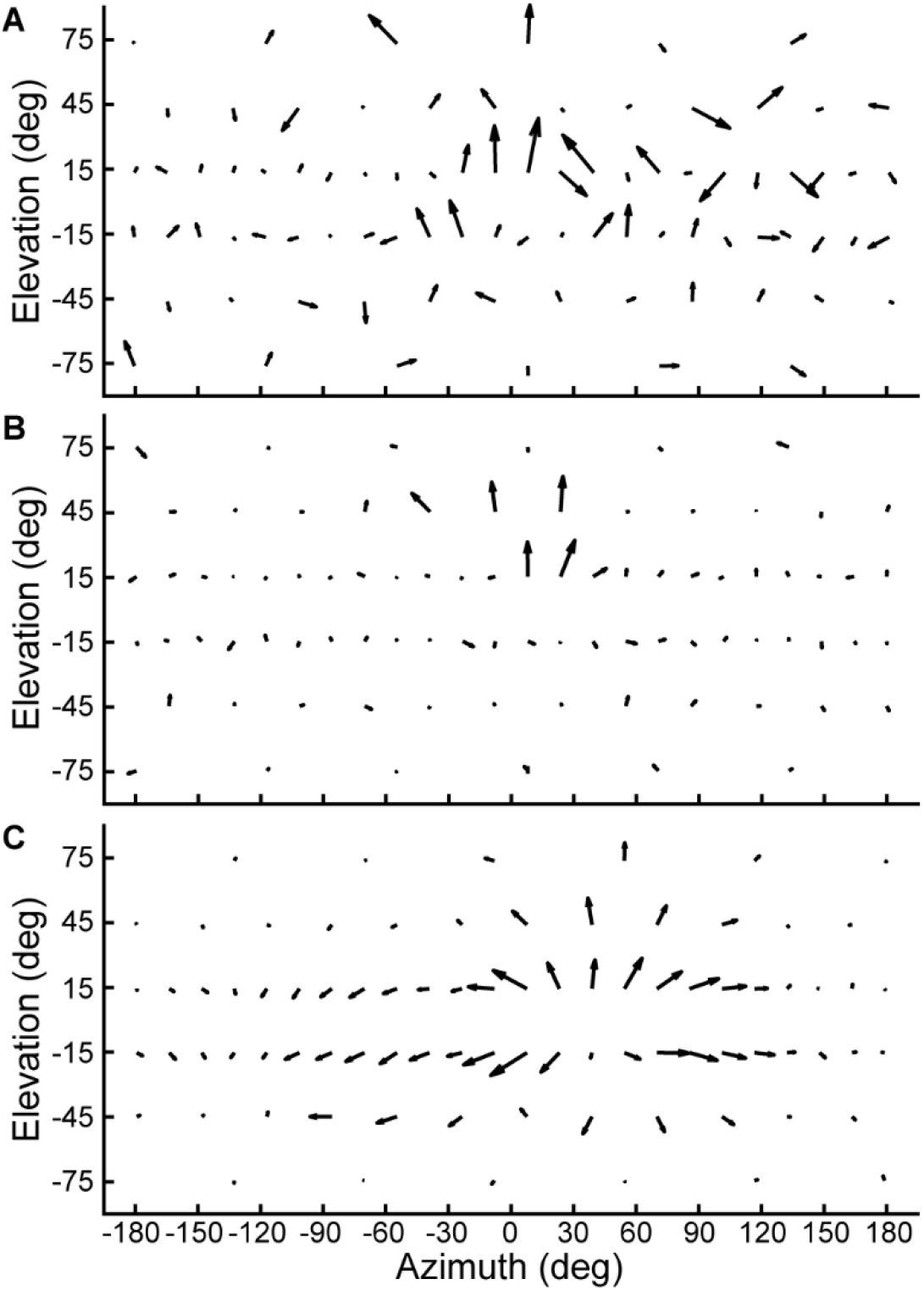
Examples of Lobula Plate Cell receptive fields not included in the KvD and phase locking analysis. **A)** Incoherent distribution of local preferred directions. **B)** Target-selective cell. **C)** Translation-sensitive cell.

**Figure S2:**
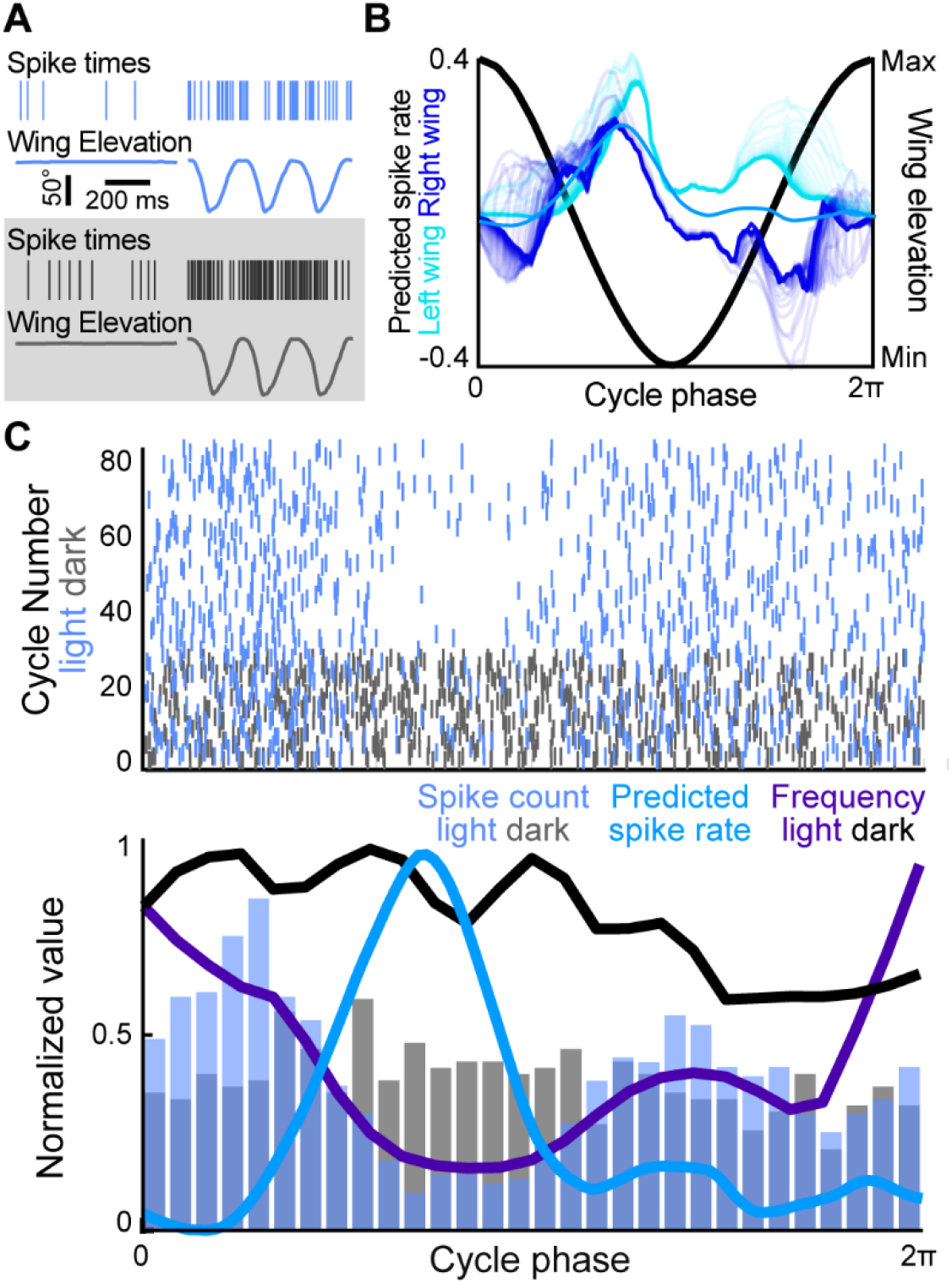
**A)** Spike times and wing elevation for a roll-sensitive LPC in the light (top) and in the dark (bottom). **B**) Predicted spike rate across all wingbeats for the left (cyan) and right (dark blue) wing trajectories. Normalized wing elevation shown in black and average predicted spike rate for both wings and all wingbeats is shown in blue. **C**) Raster plot of spike times relative to wingbeat phase in the light shown in light blue and in the dark shown in grey (top). Predicted spike response (blue line; see panel B), binned spike counts (light blue & grey histogram), and average instantaneous spike rate in the light and dark (purple and black lines). All data were normalized to the duration of the wingbeat cycle. The expected response based solely on wing movement did not predict the measured spiking activity

**Figure S3:**
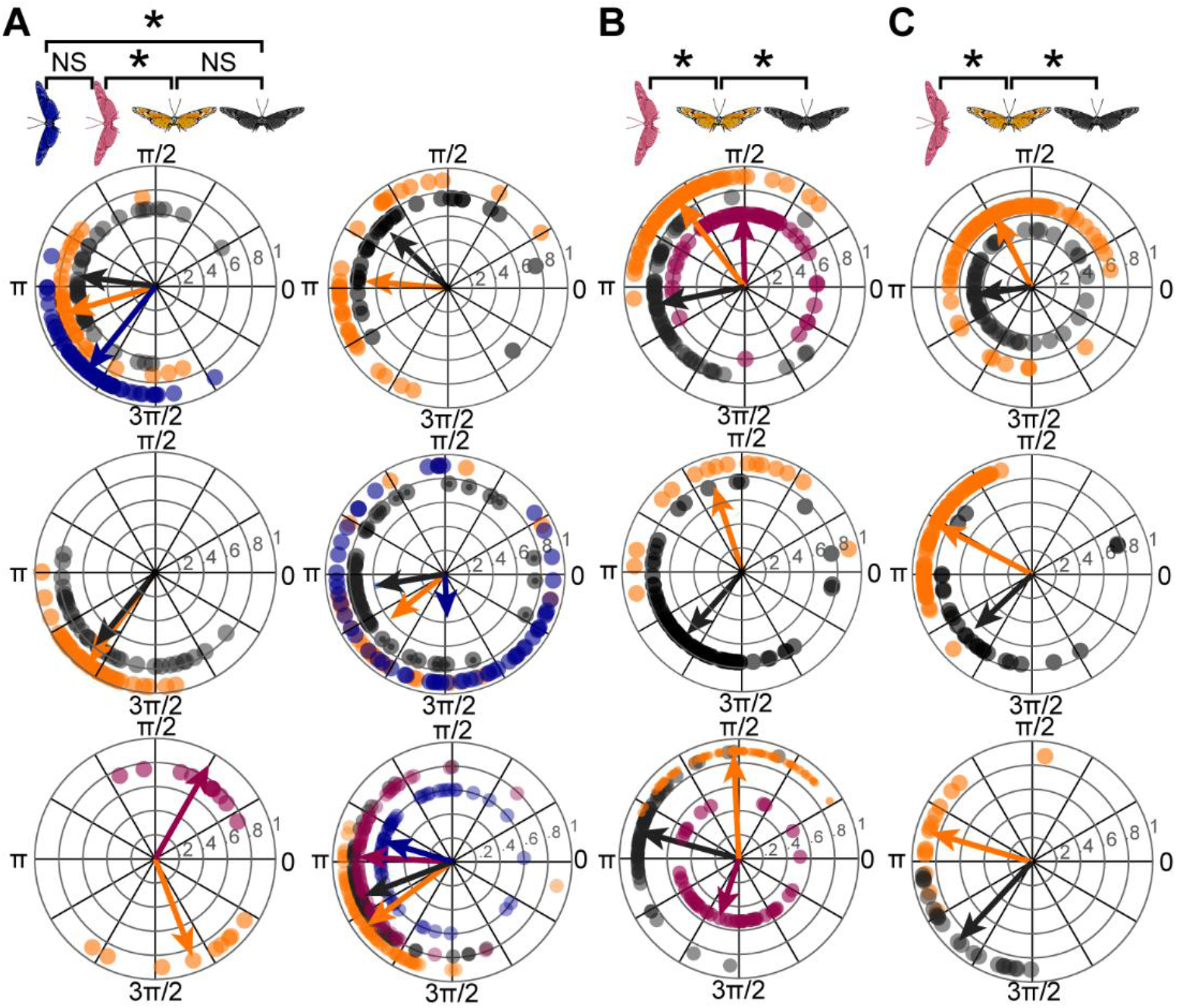
Polar plots for yaw (A), pitch (B), and roll (C) cells recorded during flapping in the light (orange) and the dark (grey) in upright and sideways (pink and purple) orientations. Body orientation significantly changed phase of activation across all cells, but light condition only affected the phase of pitch and roll cells.

**Figure S4:**
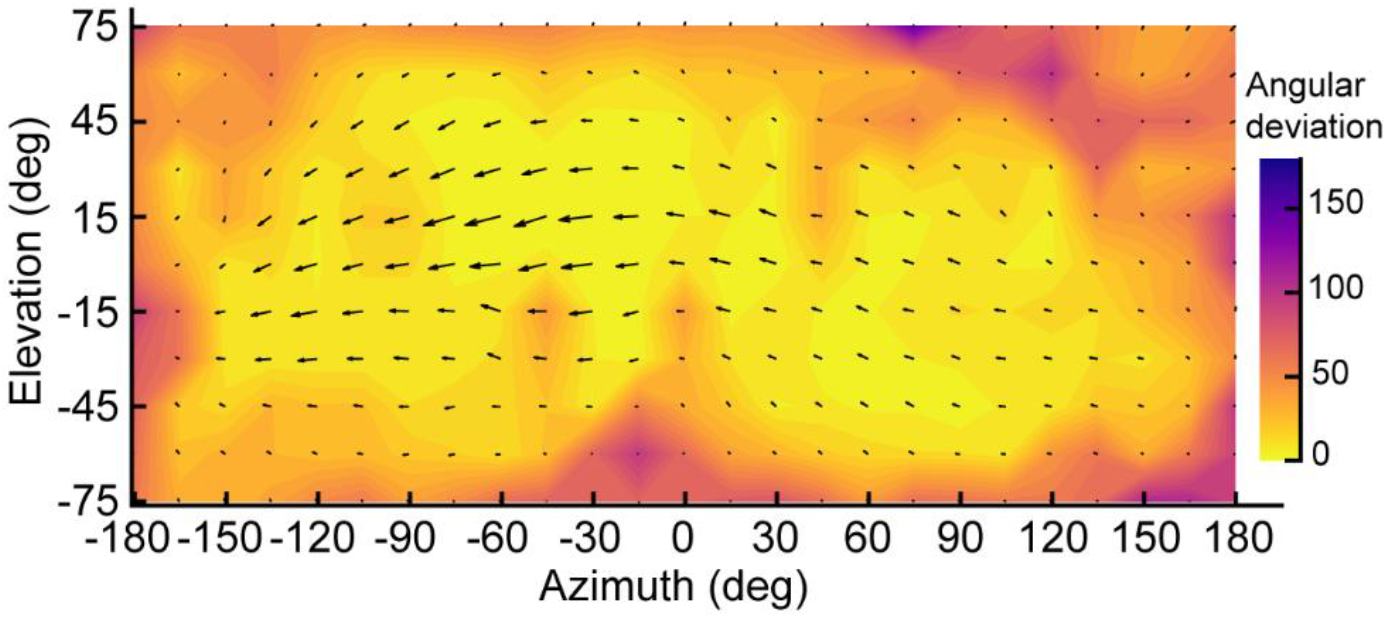
Interpolated receptive field map averaged across five trials. The heatmap shows that angular variance between trials correlates with the local motion sensitivity (vector length).

**Figure S5:**
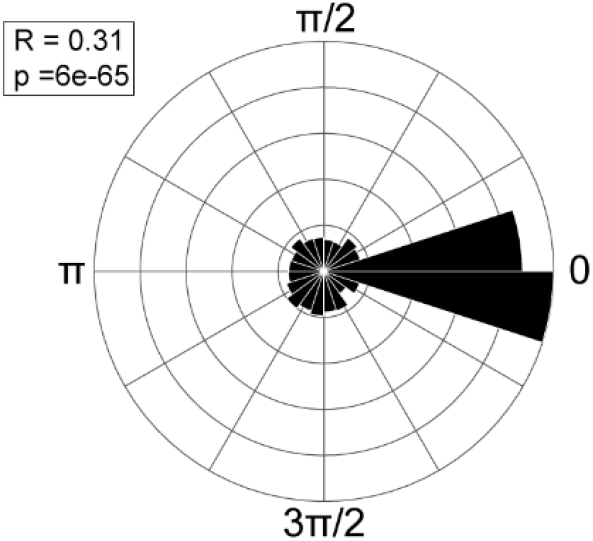
Polar plot showing significant phase locking of spikes relative to the first wingbeat cycle across all LPCs and wingbeats.

**Figure S6:**
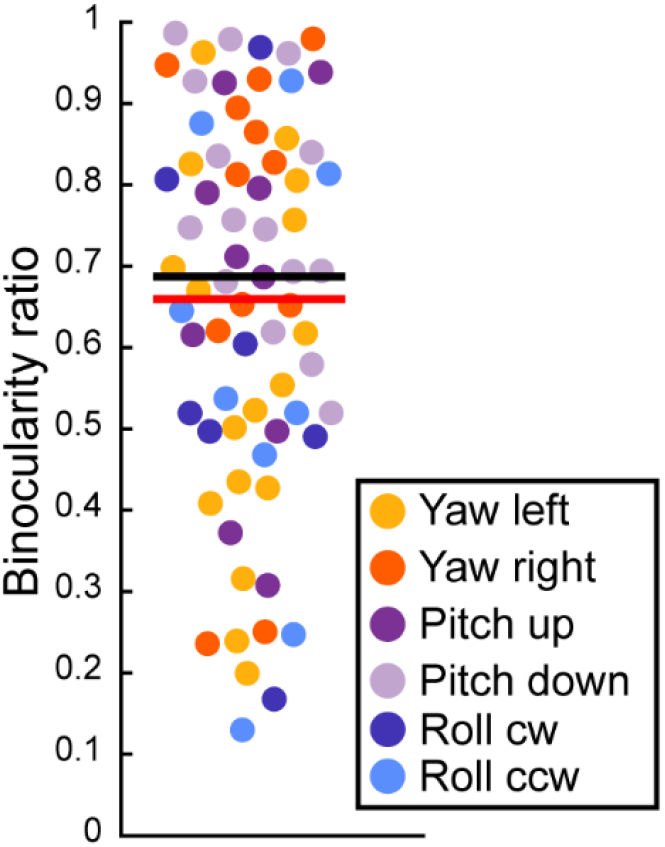
Binocularity ratios of recorded LPCs. Values closer to 1 indicate higher binocularity. 0 represents monocular responses.

**Figure S7:**
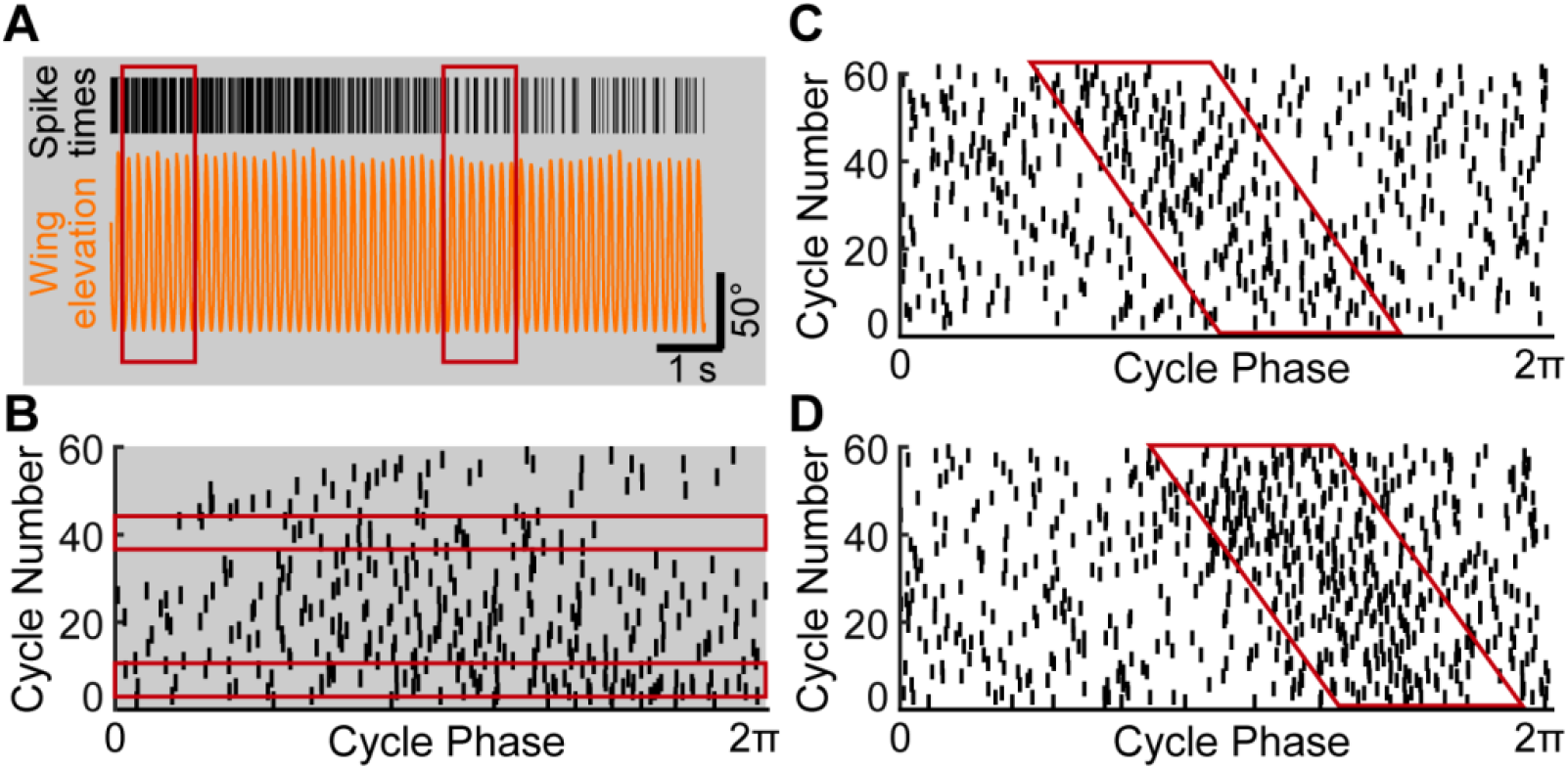
**A)** Example flapping bout in the dark for yaw cell with spike times shown in black and wing elevation in orange. **B)** Raster plot of spike times relative to wingbeat phase. Red boxes show example phase jumps corresponding to panel A and Video S1. **C**) Example raster plot of yaw cell in the light showing phase shift over time in a rotated animal. **D**) Example raster plot of pitch cell in the light showing phase shift over time in a rotated animal.

**Table S1.**
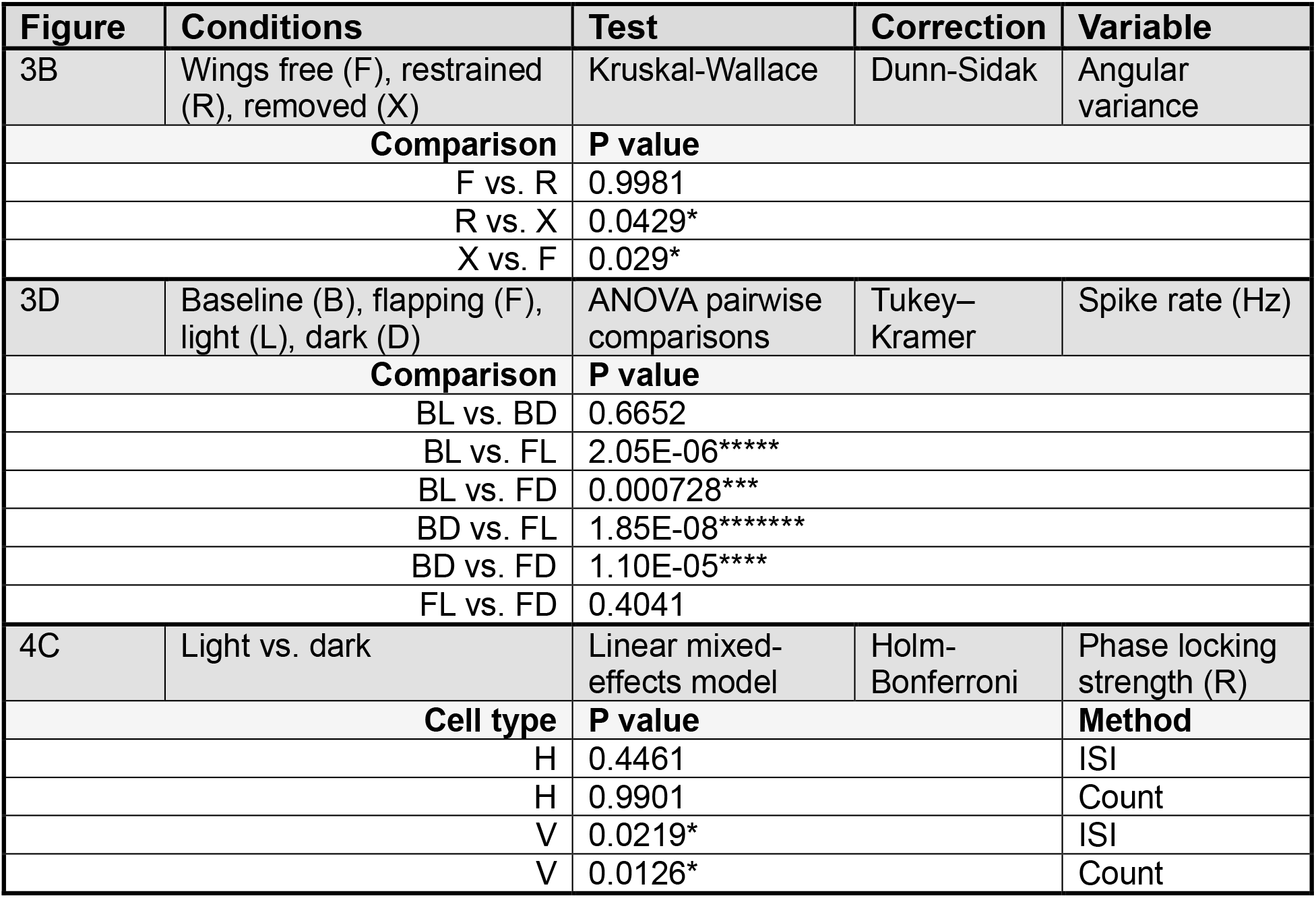
Summary of statistics.

**Video S1:** Example yaw cell phase locking in the dark at different phases with no obvious kinematic differences.

## References

1. Horrocks, E.A.B., Mareschal, I., and Saleem, A.B. Walking humans and running mice: perception and neural encoding of optic flow during self-motion. Philos Trans R Soc Lond B Biol Sci 378, 20210450. 10.1098/rstb.2021.0450.

2. Wylie, D.R. (2013). Processing of visual signals related to self-motion in the cerebellum of pigeons. Front Behav Neurosci 7, 4. 10.3389/fnbeh.2013.00004.

3. Egelhaaf, M. (2023). Optic flow based spatial vision in insects. J Comp Physiol A 209, 541–561. 10.1007/s00359-022-01610-w.

4. Krapp, H.G. (2014). Optic Flow Processing. In Encyclopedia of Computational Neuroscience (Springer, New York, NY), pp. 1–22. 10.1007/978-1-4614-7320-6_332-1.

5. Taylor, G.K., and Krapp, H.G. (2007). Sensory systems and flight stability: what do insects measure and why? Advances in insect physiology 34, 231–316.

6. Hardcastle, B.J., and Krapp, H.G. (2016). Evolution of Biological Image Stabilization. Curr Biol 26, R1010–R1021. 10.1016/j.cub.2016.08.059.

7. Krapp, H.G. (2015). How a fly escapes the reflex trap. Nat Neurosci 18, 1192–1194. 10.1038/nn.4097.

8. Wolpert, D.M., Miall, R.C., and Kawato, M. (1998). Internal models in the cerebellum. Trends Cogn Sci 2, 338–347. 10.1016/s1364-6613(98)01221-2.

9. Kim, A.J., Fenk, L.M., Lyu, C., and Maimon, G. (2017). Quantitative Predictions Orchestrate Visual Signaling in Drosophila. Cell 168, 280–294.e12. 10.1016/j.cell.2016.12.005.

10. Kim, A.J., Fitzgerald, J.K., and Maimon, G. (2015). Cellular evidence for efference copy in Drosophila visuomotor processing. Nat Neurosci 18, 1247–1255. 10.1038/nn.4083.

11. Fenk, L.M., Kim, A.J., and Maimon, G. (2021). Suppression of motion vision during course-changing, but not course-stabilizing, navigational turns. Current Biology 31, 4608–4619. 10.1016/j.cub.2021.09.068.

12. Falt, T., Ammer, G., Serbe-Kamp, É., Kroell, L.M., Friedrich, A.B., Guerreiro-Mota, S., Šimsová, E., and Fenk, L.M. (2026). Vectorial efference copy and visuomotor transformation through gap junctions. Preprint at bioRxiv, 10.64898/2026.08.11.744090 https://doi.org/10.64898/2026.08.11.744090.

13. Krapp, H.G., and Hengstenberg, R. (1996). Estimation of self-motion by optic flow processing in single visual interneurons. Nature 384, 463–466. 10.1038/384463a0.

14. Keesey, I.W., Knaden, M., and Hansson, B.S. (2015). Olfactory Specialization in Drosophila suzukii Supports an Ecological Shift in Host Preference from Rotten to Fresh Fruit. J Chem Ecol 41, 121–128. 10.1007/s10886-015-0544-3.

15. Jernigan, C.M., Halby, R., Gerkin, R.C., Sinakevitch, I., Locatelli, F., and Smith, B.H. (2020). Experience-dependent tuning of early olfactory processing in the adult honey bee, Apis mellifera. J Exp Biol 223, jeb206748. 10.1242/jeb.206748.

16. Dong, L., Hormigo, R., Barnett, J.M., Greppi, C., and Duvall, L.B. (2025). Time-of-day modulation in mosquito response persistence to carbon dioxide is controlled by Pigment-Dispersing Factor. Proceedings of the National Academy of Sciences 122, e2520826122. 10.1073/pnas.2520826122.

17. Rien, D., Kern, R., and Kutz, R. (2013). Octopaminergic modulation of a fly visual motion-sensitive neuron during stimulation with naturalistic optic flow. Frontiers in behavioral neuroscience 7. 10.3389/fnbeh.2013.00155.

18. Longden, K.D., Muzzu, T., Cook, D.J., Schultz, S.R., and Krapp, H.G. (2014). Nutritional State Modulates the Neural Processing of Visual Motion. Current Biology 24, 890–895. 10.1016/j.cub.2014.03.005.

19. Longden, K.D., and Krapp, H.G. (2010). Octopaminergic Modulation of Temporal Frequency Coding in an Identified Optic Flow-Processing Interneuron. Front. Syst. Neurosci. 4. 10.3389/fnsys.2010.00153.

20. Maimon, G., Straw, A.D., and Dickinson, M.H. (2010). Active flight increases the gain of visual motion processing in Drosophila. Nat Neurosci 13, 393–399. 10.1038/nn.2492.

21. Suver, M.P., Mamiya, A., and Dickinson, M.H. (2012). Octopamine neurons mediate flight-induced modulation of visual processing in Drosophila. Curr Biol 22, 2294–2302. 10.1016/j.cub.2012.10.034.

22. Jung, S.N., Borst, A., and Haag, J. (2011). Flight activity alters velocity tuning of fly motion-sensitive neurons. J Neurosci 31, 9231–9237. 10.1523/JNEUROSCI.1138-11.2011.

23. Fayyazuddin, A., and Dickinson, M.H. (1996). Haltere afferents provide direct, electrotonic input to a steering motor neuron in the blowfly, Calliphora. J Neurosci 16, 5225–5232. 10.1523/JNEUROSCI.16-16-05225.1996.

24. Trischler, C., Kern, R., and Egelhaaf, M. (2010). Chasing behaviour and optomotor following in free-flying male blowflies: flight performance and interactions of the underlying control systems. Front. Behav. Neurosci. 4. 10.3389/fnbeh.2010.00020.

25. Fabian, S.T., Yarger, A.M., Chen, S.-L., and Lin, H.-T. (in press). The aerial combat strategy of dragonflies. J. R. Soc. Interface.

26. Guerra, P.A., Gegear, R.J., and Reppert, S.M. (2014). A magnetic compass aids monarch butterfly migration. Nat Commun 5, 4164. 10.1038/ncomms5164.

27. Perez, S.M., Taylor, O.R., and Jander, R. (1997). A sun compass in monarch butterflies. Nature 387, 29–29. 10.1038/387029a0.

28. Froy, O., Gotter, A.L., Casselman, A.L., and Reppert, S.M. (2003). Illuminating the Circadian Clock in Monarch Butterfly Migration. Science 300, 1303–1305. 10.1126/science.1084874.

29. Sherman, A., and Dickinson, M.H. (2004). Summation of visual and mechanosensory feedback in Drosophilaflight control. J Exp Biol 207, 133–142. 10.1242/jeb.00731.

30. Bender, J.A., and Dickinson, M.H. (2006). A comparison of visual and haltere-mediated feedback in the control of body saccades in Drosophila melanogaster. Journal of Experimental Biology 209, 4597–4606. 10.1242/jeb.02583.

31. Dahake, A., Stöckl, A.L., Foster, J.J., Sane, S.P., and Kelber, A. (2018). The roles of vision and antennal mechanoreception in hawkmoth flight control. eLife 7, e37606. 10.7554/eLife.37606.

32. Huston, S.J., and Krapp, H.G. (2009). Nonlinear Integration of Visual and Haltere Inputs in Fly Neck Motor Neurons. J. Neurosci. 29, 13097–13105. 10.1523/JNEUROSCI.2915-09.2009.

33. Allibert, G., Courtial, E., and Chaumette, F. (2010). Visual Servoing via Nonlinear Predictive Control. In Visual Servoing via Advanced Numerical Methods Lecture Notes in Control and Information Sciences., G. Chesi and K. Hashimoto, eds. (Springer London), pp. 375–393. 10.1007/978-1-84996-089-2_20.

34. Fujiwara, T., Cruz, T.L., Bohnslav, J.P., and Chiappe, M.E. (2017). A faithful internal representation of walking movements in the Drosophila visual system. Nat Neurosci 20, 72–81. 10.1038/nn.4435.

35. Weber, A.I., Daniel, T.L., and Brunton, B.W. (2021). Wing structure and neural encoding jointly determine sensing strategies in insect flight. PLoS Comput Biol 17, e1009195. 10.1371/journal.pcbi.1009195.

36. Aiello, B.R., Stanchak, K.E., Weber, A.I., Deora, T., Sponberg, S., and Brunton, B.W. (2021). Spatial distribution of campaniform sensilla mechanosensors on wings: form, function, and phylogeny. Current Opinion in Insect Science 48, 8–17. 10.1016/j.cois.2021.06.002.

37. Koenderink, J.J., and van Doorn, A.J. (1987). Facts on optic flow. Biol Cybern 56, 247–254. 10.1007/BF00365219.

38. Red-green opponency in the long visual fibre photoreceptors of brushfoot butterflies (Nymphalidae) | Proceedings B | The Royal Society https://royalsocietypublishing.org/rspb/article/288/1961/20211560/79193/Red-green-opponency-in-the-long-visual-fibre.

